# Artificial feeding of invasive grey squirrels (*Sciurus carolinensis*) at urban parks: a social norms approach to understand its drivers and to guide behavioral interventions

**DOI:** 10.1101/2020.07.15.205260

**Authors:** Jacopo Cerri, Elena Martinelli, Sandro Bertolino

## Abstract

Artificial wildlife feeding might contribute to the successful establishment of some invasive alien species, like the Eastern grey squirrel (Sciurus carolinensis) in Europe and the UK. Reducing squirrel feeding at urban parks can be important to reduce squirrel populations while avoiding social conflicts. From April to October 2018, we conducted interviews and administered factorial surveys to two samples of visitors at the Valentino urban park in Torino (Italy). We established whether squirrel feeding can be regarded as an independent or an interdependent behavior, by eliciting visitors’ moral beliefs, and their empirical and normative expectations.

Most respondents did not regard artificial feeding as something neither intrinsically positive nor negative. Satisfaction, the need for a connection with nature and the presence of children were its main causes. On the other hand, feeding squirrels was deemed to be potentially dangerous for squirrel health, and leading to high squirrel densities and to confident squirrels.

Factorial surveys indicated that empirical expectations about the behavior of other visitors, altogether with their past behavior, lead visitors to feed squirrels, while the availability of suboptimal food types (e.g. sweets), was deemed to made people less prone to forage squirrels. Our findings indicate that supplementary feeding is an interdependent behavior, governed by descriptive norms. It might be reduced by means of informative panels containing descriptive information about the fact that most visitors indeed do not feed squirrels, as well as by panels emphasizing the potential damage that squirrels could suffer from eating suboptimal food provided by humans.

## Introduction

The artificial feeding of wild animals is one of the most important human-wildlife interactions in urbanized ecosystems, either in a positive and in a negative way (Dubois and Fraser 2013). Available evidence suggests that it can strengthen connection with nature, inform people about conservation and foster environmental stewardship (Cox and Gaston 2016; Larson et al. 2016). However, artificial feeding can also have negative consequences, both for people and wildlife. For example, it can increase the risk of disease transmission (Hwang et al. 2018; Murray et al. 2016), can make wildlife confident towards humans and can disrupt ecological balance among wildlife populations (Galbraith et al. 2015; Jones and Raynolds 2008), either by reducing natural mortality and by affecting dispersal (Richardson et al. 2015; Starkey 2019).

This latter point can also have major implications for invasive alien species (IAS), by making easier for them to settle down in urbanized areas and to establish viable populations. For example, the Eastern grey squirrel (*Sciurus carolinensis*, hereafter grey squirrel) successfully colonized Italy, Ireland and the UK, in many cases after having established viable populations at large urban parks, where the species is regularly foraged by visitors (Bertolino 2009, Bertolino et al. 2014). This species is widespread in Central and Northern Italy (Loy et al. 2019) and is considered invasive because it causes the extinction of the native Eurasian red squirrel (*Sciurus vulgaris*) (Gurnell et al. 2004). Since it is expected that grey squirrels could further spread in Italy and in Europe (Di Febbraro et al. 2016, 2019), the species should be eradicated or at least numerically reduced in many areas (Bertolino et al. 2014). The connection that could be established between squirrels and visitors in urban areas can make any management intervention to limit exotic squirrels difficult to implement for the opposition of groups of visitors (Bertolino et al. 2015; Lioy et al. 2016). Urban parks are therefore strongholds, where grey squirrels are an iconic species, beloved by visitors who maintain their densities high through supplementary feeding (Merrick et al. 2016), often opposing to their control. Even if conservationists were able to remove grey squirrels from woodlands and croplands, at the large geographical scale, a source-sink system is likely to appear, with urban parks acting as a source of squirrels who will disperse around, undermining eradication efforts. Reducing the supplementary feeding of squirrels by visitors could be important to avoid this scenario. Reducing squirrel numbers at urban parks could also decrease their interactions with visitors, making in turn them less iconic in the long term.

Behavioral interventions, such as informative panels providing visitors with descriptive or normative messages, might be a valuable tool to reduce supplementary feeding of wildlife and to reduce disturbance to wildlife (Mackay et al. 2018; Marion et al. 2008). Normative messages seems to be particularly promising, because social norms act over human behavior without requiring attitude change, and because behavioral change can in turn make them more powerful through time, creating positive feedbacks which increase their effectiveness in the long term (Farrow et al. 2017; Schultz et al. 2018). However, behavioral interventions should be carefully designed, according to the nature of the behavior they target (Hauser et al. 2019). For example, panels adopting normative messages to target highly individual behaviors will not be effective. Panels providing the wrong information (e.g. about norm defection) might also backfire, going into the wrong direction and triggering norm defection and the undesirable behavior they should have discouraged (Bicchieri and Dimant 2019).

Bicchieri et al. (2016) provided an operational taxonomy to evaluate collective human behaviors, ranging from highly independent to highly interdependent ones, such as social norms. The taxonomy classifies behaviors based on five components: *i*) practical usefulness, *ii*) moral beliefs, *iii*) empirical expectations, *iv*) normative expectations and *v*) sanctions and rewards. In turn, these five components identify two types of independent behaviors, customs and moral norms, and two types of interdependent human behaviors, namely descriptive and social norms. Independent behaviors are carried out regardless of what others do: customs occur because there are practical reasons for them to exist (e.g. using an umbrella when it rains), and moral norms because people hold strong beliefs about the morality of a certain behavior (e.g. animal right activism). On the other hand, interdependent behaviors are conditional on our expectations about others. Descriptive norms govern behaviors through empirical expectations alone: we believe a certain action to be regularly done by people in our reference network, then we do it as well (e.g. free-riding natural resources). Finally, social norms rely on empirical and normative expectations, altogether with informal sanctions/rewards associated to their defection/enforcement. Human behaviors are subjected to social norms, because people expect others to engage in these behaviors (empirical expectations), to have expectations as well (normative expectations) and to believe in rewards associated with endorsement (e.g. approval, reputation) and sanctions associated with defection (e.g. disapproval) (Fig. 1).

**Figure 1.**
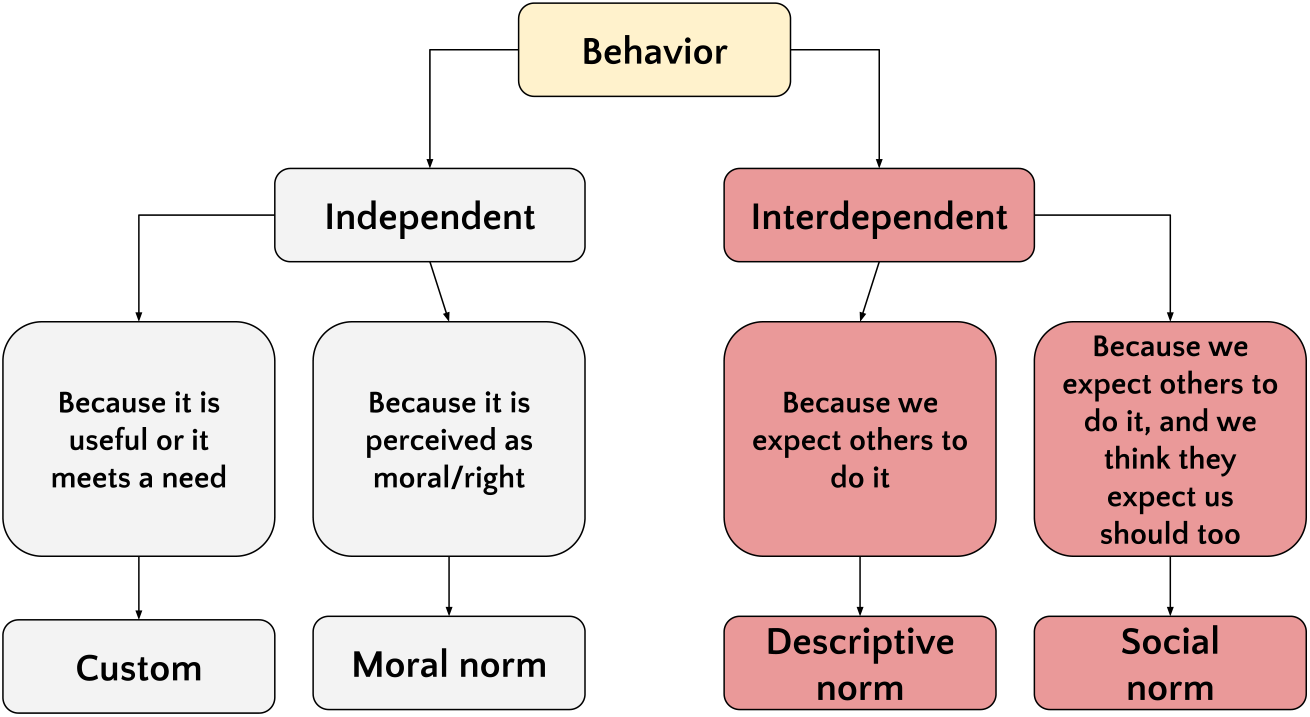
Taxonomy of collective behaviors according to Bicchieri et al. (2016). The scheme was inspired by Bicchieri and Noah (2017).

This classification can be extremely useful to classify human-wildlife interactions, such as artificial feeding, and to suggest practical behavioral interventions targeting them. For example, changing empirical expectations through informative panels is fundamental for descriptive norms, but it might be ineffective for moral norms and it should be coupled with the identification of role models and trend-setters for social norms (Bicchieri et al. 2016).

In this study, by combining qualitative interviews with quantitative factorial surveys, we aim to show how this classification can identify the main drivers of supplementary feeding of grey squirrels at urban parks, and how it can inform practical interventions to reduce it.

## Methods

### Study area

The study was carried out at the Valentino urban park, in the city of Torino (Italy, 45°03’24.3“N 7°41’18.2”E). The park covers a surface of 42 hectares, over the western bank of the Po river, and its land cover includes a mixture of native and ornamental trees and shrubs, patches of mixed woodlots, meadows and a botanical garden. The park hosts a population of grey squirrels, which were introduced in 1948 near Stupinigi, at approximately 10 km from the study area (Bertolino 2009). The population density inside the Valentino park is not known; however, the species can reach in urban parks densities up to 20 individuals/ha (Venturini et al. 2005; Merrick et al. 2016). Nowadays, it is far common for visitors to encounter grey squirrels, when visiting the park, and they became an iconic species, which is regularly encountered and foraged by visitors. The red squirrel, present in the park in the past, has not been reported for many years.

### Qualitative interviews

Between March and April 2018, we carried out structured qualitative interviews with a sample of visitors (n = 32) at the park. Qualitative methods are useful to obtain insights about human-wildlife relationships, when little or no preliminary evidence is available about them and when researchers want to acquire some unexpected element that might then be included in structured surveys (Drury et al. 2011; Rust et al. 2017).

In our case we asked respondents about the presence, within their reference network of figures who might act as role models, of people who might oppose or support supplementary feeding of squirrels at the park. For each one of the two groups, those who approved and those who disapproved, we asked who they were (e.g. relatives, friends) as well as about empirical expectations, normative expectations and beliefs about norm endorsement. Then we asked participants for their moral beliefs about artificial feeding, by providing them with a 5-points bipolar scale ranging from “Completely wrong” to “Totally right”, and by asking them to use it to rate their beliefs. We asked participants also to tell us the potential benefits and risks, of engaging in artificial feeding while visiting the park, and how hard they deemed to be to quit feeding squirrels for an habitué. Finally, to measure norm endorsement, we asked people to imagine meeting someone feeding squirrels when visiting the park, and if they would disapprove or not this behavior and whether they would reprove the visitor in public. Finally, after having ensured the confidentiality of our findings, we asked respondents to provide us with some of their personal details (e.g. age, sex, education; Table 1). Interviews lasted approximately 10 minutes, we used an audio recorder to record answers and then we transcribed them. Answers were coded manually, and we used findings from the interviews to design the successive factorial survey experiment.

**Table 1:** Structure of interviews (questions asked during the interview) and factorial surveys (factors and their levels)

| Qualitative interviews<br>Construct | Question |
| --- | --- |
| Reference network: people who would disapprove artificial feeding of squirrels at urban parks | If you came here at the Valentino park, for example during the next two weeks, some people might believe you should NOT feed the squirrels. Could you please tell us if some of these people come to your mind, and who they are |
| Empirical expectations | Do you believe these people would avoid feeding squirrels, if they visited the park? |
| Normative expectations | Do you believe these people would deem inappropriate for you to feed squirrels at the park? |
| Norm endorsement (reference network) | In your opinion, if these people noticed a visitor feeding a squirrel, while visiting the park, would they do something about it? What ? |
| Reference network: people who would approve artificial feeding of squirrels at urban parks | If you came here at the Valentino park, for example during the next two weeks, some people might believe you should NOT feed the squirrels. Could you please tell us if some of these people come to your mind, and who they are? |
| Empirical expectations | Do you believe these people would feed squirrels, if they visited the park? |
| Normative expectations | Do you believe these people would approve you, in case you feed the squirrels at the park? |
| Moral beliefs | Look at the scoring card you received. Could you please tell us, based on the scores reported on it, how wrong or right you believe squirrel feeding to be? |
| Practical reasons for artificial feeding | Based on what you know and heard about it, could you tell us which are the positive aspects of feeding squirrels at the park? |
| Negative consequences of artificial feeding | And could you also tell us which might be the negative consequences of feeding squirrels at the park? |
| Perceived difficulty at abandoning the target behavior | In your opinion, how hard would be, for a visitor who regularly feeds squirrels, to stop doing it? Why? |
| Norm endorsement (respondent): private and public disapproval? | Now, please imagine this situation: you are walking at the park and you notice another visitor who is feeding a squirrel. Would you disapprove such behavior, even without saying anything to the visitor? Would you do something about it, like reproving the visitor? |
| Age | Could you please tell us your age? |
| Sex (reported by interviewer) | - |
| Level of education | Could you please tell us which is your level of education? Do you have a degree? |
| Geographical origin | Where are you born in? |
| Residence | Do you live in Torino? |
| Parenthood | Do you have any son or daughter? |
| Dog ownership | Do you possess any dog? |
| Frequency visit | On average, how often do you visit the Valentino park? |
| Factorial surveys |  |
| Factor | Levels |
| Food type (vignette) | The visitor has some nuts, as a snack<br>The visitor has some cookies, as a snack |
| Presence of children (vignette) | The visitor is walking at the park with his/her daughter, who is 5 years old<br>The visitor is walking alone |
| Past behavior (vignette) | The visitor never fed squirrels at the park<br>In the past, the visitor fed the squirrels sometimes |
| Visibility (vignette) | On that moment, other people at the park could see the visitor<br>On that moment, the visitor cannot be seen by any other people |
| Other visitors feeding squirrels (vignette) | During that visit at the park, he/she did not see anyone feeding squirrels<br>During that visit at the park, he/she saw some other visitors who were feeding squirrels |
| Season (vignette) | It is winter and it's cold<br>It is summer and there is a nice weather |
| Empirical expectations (vignette) | The visitor believes that someone important to she/him, would not feed the squirrel on that occasion<br>The visitor believes that someone important to she/him, would feed the squirrel on that occasion |
| Response variable (vignette) | In your opinion, would the visitor feed the squirrel on that occasion? (Yes; No) |
| Residency (respondent) | Do you live in Torino? (Yes; No) |
| Age (respondent) | How old are you? (18-25 years; 26-35 years; 36-45 years; 46-60 years; more than 60 years) |
| Sex | You are... (Man; Woman; I would rather not to say it) |
| Education | Level of education... (elementary; secondary school; high school; university degree or higher) |
| Frequency of visit | How often do you visit this park? (First time; Not every year; Multiple times every year; Multiple times every month; Multiple times every week) |

### Factorial surveys

Between May and September 2019, we also designed and carried out a factorial survey experiment with a sample of visitors (n = 32). Factorial surveys are a particular form of conjoint evaluation method, where each respondent is asked to evaluate a sample of hypothetical situations, termed “vignettes”. Vignettes are characterized by a series of attributes describing the situation, which are combined through a factorial design (Auspurg and Hinz 2014), just like an experiment. In our case, we created 256 vignettes depicting a hypothetical situation where a visitor was strolling through the park, with some food in his hand, and was approached by a confident grey squirrel. For each vignette we described the type of food held by the visitor (nuts or a cookie), whether the visitor was in company of a child, whether he was walking the dog, the time of the year (winter or summer), if the visitor was visible to others in that moment and whether he/she had seen some visitors feeding squirrels previously. Finally, we characterized the vignettes in terms of empirical expectations, by specifying whether the visitor believed other visitors in his/her reference network would have fed the squirrel, on that occasion. Given the conditions described in the vignette, we asked respondents to specify whether they believed the visitor in the vignette would forage the squirrel or not. A vignette appeared as it follows:

> A visitor is walking at the Valentino urban park, when he/she notices a grey squirrel approaching. The visitor has some nuts, as a snack, and believes that someone important to her/him, would not feed the squirrel on that occasion. During that visit at the park, he/she saw some other visitors who were feeding squirrels and, on that moment, the visitor cannot be seen by any other people. In the past, the visitor fed the squirrels sometimes. It is summer, there is a nice weather and the visitor is walking alone.
>
> In your opinion, would the visitor feed the squirrel on that occasion? (Yes/No)

Our factorial design aimed to test whether empirical expectations, situational factors, and previous experience, can be important in the decision to engage in supplementary feeding. We carried out hierarchical Bayesian Generalized Linear Modeling (Gelman et al. 2013), by specifying a binomial structure of the error with a logarithmic link function, and by specifying a random intercept term to account for heterogeneity between respondents. As visitors from Torino could have more salient beliefs about park usage than tourists, and different vignette evaluation, we introduced a dichotomous second-order variable distinguishing residents from non-residents among respondents, and we controlled for it in the model. We used uninformative uniform priors for model parameters, and we fit 4 Markov chains with 5000 iterations each. Model selection was based on the widely applicable information criterion (WAIC) and k-fold cross validation with 10 folds. We checked for MCMC convergence and we carried out posterior predictive checks from the posterior distribution of model parameters. Bayesian hierarchical modeling was carried out with STAN (Gelman et al. 2015), through the statistical package ‘brms’ (Bürkner 2018) of the software R (R Core Team 2019).

## Results

### Qualitative interviews

Overall, 50% of respondents were men, age was 44.1 ± 19.7 years (mean ± sd), 75.0% of them were from Torino and their level of education was high, with 55.6% of respondents having a university degree. Most respondents also visited the park with a frequency between one time per week and everyday (68.5%).

About 30.6% of respondents reported to know someone who would disapprove the supplementary feeding of squirrels at the park, mostly relatives (45.5%), friends (45.5%) or other visitors (18.2%). Overall 90.1% of these visitors had empirical expectations about these persons, believing them not to feed squirrels, when visiting the park. 72.7% of these visitors also had normative expectations, believing these people would disapprove them in case they fed squirrels. Only few of them believed these people would reprove squirrel feeding in public (18.1%), when visiting the park. About 27.8% of respondents, on the contrary, reported to knew someone who would openly approve the supplementary feeding of squirrels, mostly relatives (45.4%) or friends (45.4%). Overall 90% of these visitors have positive empirical expectations about these people and 80% of them had normative expectations.

Moral beliefs were relatively neutral, as 48.6 % of interviewed did not regarded feeding as something intrinsically positive or negative. The main reasons why they believed visitors to feed squirrels were psychological pleasure (38.9%), the need for a connection with nature and wildlife (38.9%) or the presence of children (27.8%). On the other hand, they believe squirrel feeding to have also potentially negative consequences, mostly their numerical increase (25.0%) and the fact that provided food might be unhealthy for squirrels (22.2%). Most respondents (71.4%) also deemed hard to quit feeding squirrels, for visitors who do so regularly, mostly because of habit. Finally, only 25% of respondents would disapprove a visitor who feed squirrels at the park, in case they saw one, and 13.7% of them would reprove it in public. However, about half of them declared they would have reproved it, in case the provided food was unhealthy for squirrels.

### Factorial surveys

Respondents who took part into the factorial survey experiment were 50.0% men and they had an heterogeneous age distribution (18-25 years = 30.0%, 26-35 years = 13.3%, 36-45 years = 26.7%, 46-60 years = 23.3%, >60 years = 6.7%) and had a high level of literacy, as 76.7% of them had a high school or a university degree. Most of them (83.3%) were resident and visited the park on a regular basis (first time = 13.3v, not every year = 3.3%, multiple times per year = 30.0%, multiple times per month =33.3%, multiple times per week = 20.0%). Model selection based on the WAIC and k-fold crossed validation retained a final model accounting for three predictors only: empirical expectations, past behavior and the type of food (Fig. 2). The final model accounted for about 31.2% of variability in respondents’ evaluation (Table 2).

**Figure 2.**
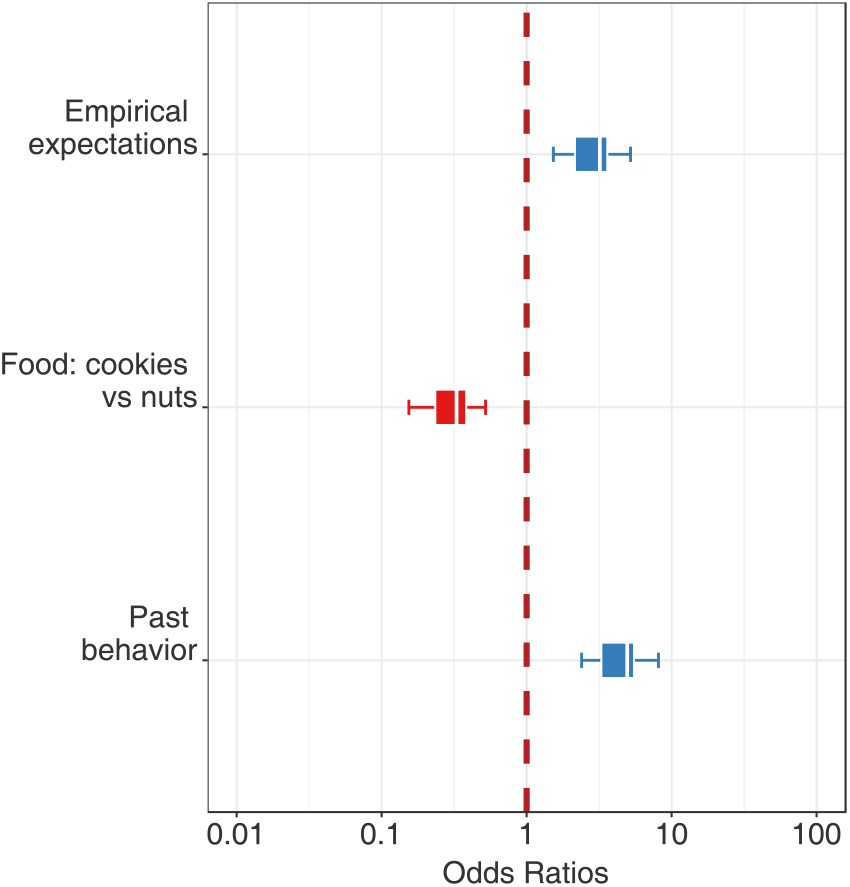
Model coefficients, odds ratios of the effect of empirical expectations, food type and past behavior of the visitor described in the vignette.

**Table 2:** Model selection: widely applicable information criterion (WAIC), k-fold cross validation with 10 folds (k-fold) and R^2^

| Model structure | WAIC | k-fold | $R^2$ |
| --- | --- | --- | --- |
| ~food type + past + empirical expectations + (1 \|*respondent*) | 267.20 | 271.08 | 0.31 |
| ~food type + past + visible + empirical expectations + (1 \|*respondent*) | 268.06 | 271.96 | 0.31 |
| ~food type + past + visible + resident + empirical expectations + (1 \|*respondent*) | 269.09 | 273.33 | 0.32 |
| ~food type + past + visible + resident + empirical expectations + visitors feeding + (1 \|*respondent*) | 268.39 | 269.03 | 0.33 |
| ~food type + past + visible + child + resident + empirical expectations + visitors feeding + (1 \|*respondent*) | 268.06 | 271.96 | 0.31 |
| ~food type + past + visible + child + resident + empirical expectations + visitors feeding + (1 \|*respondent*) | 269.32 | 266.98 | 0.34 |
| ~food type + past + visible + child + resident + empirical expectations + visitors feeding + season + (1 \|*respondent*) | 271.35 | 269.73 | 0.34 |

**Table 3:**
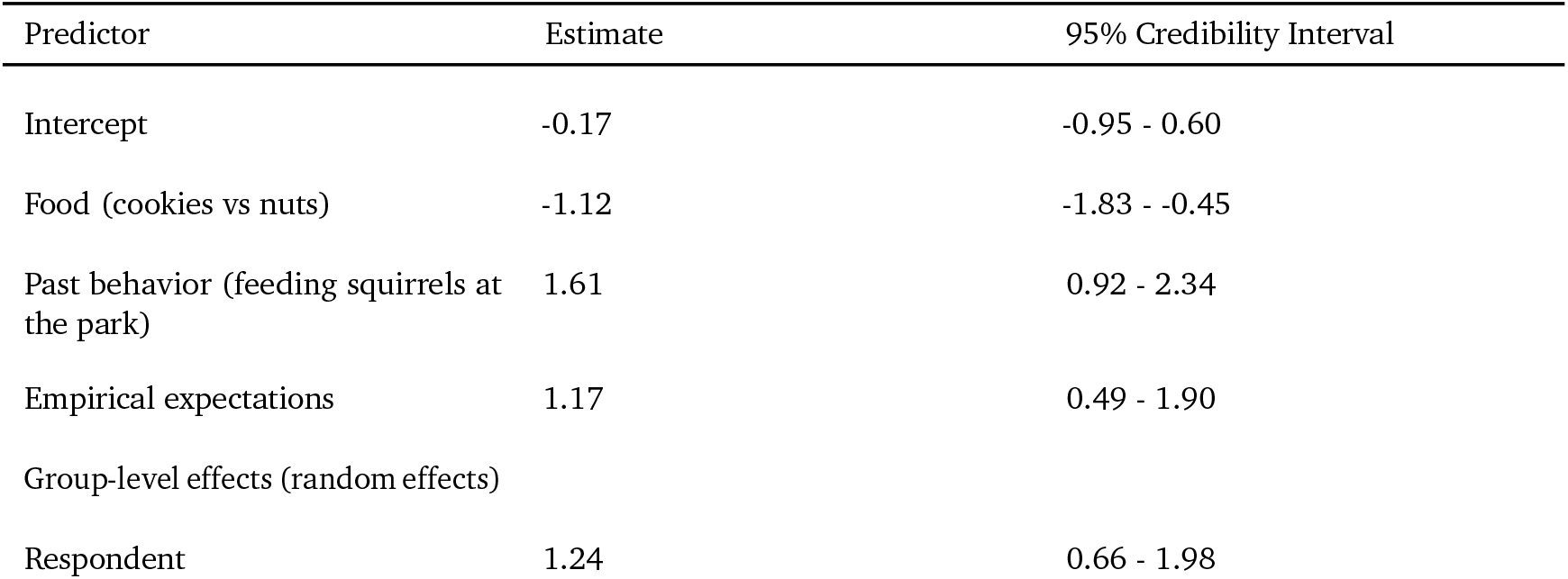
Output of hierarchical Bayesian Generalized Linear Model

Past behavior was deemed to be the most important variable affecting the probability of feeding squirrels: visitors who had already foraged squirrels in the past were regarded as 5 times more likely to do it again, in the hypothetical scenario. Empirical expectations also had a positive effect: visitors who believed other visitors to feed squirrels at the park, were perceived to be 3.2 times more likely to forage squirrels themselves. Food quality was also important: when the visitor in the vignette held a cookie, rather than nuts or dry fruits, he/she was perceived as 0.3 times less likely to give it to the squirrel as a food (Fig. 3).

**Figure 3.**
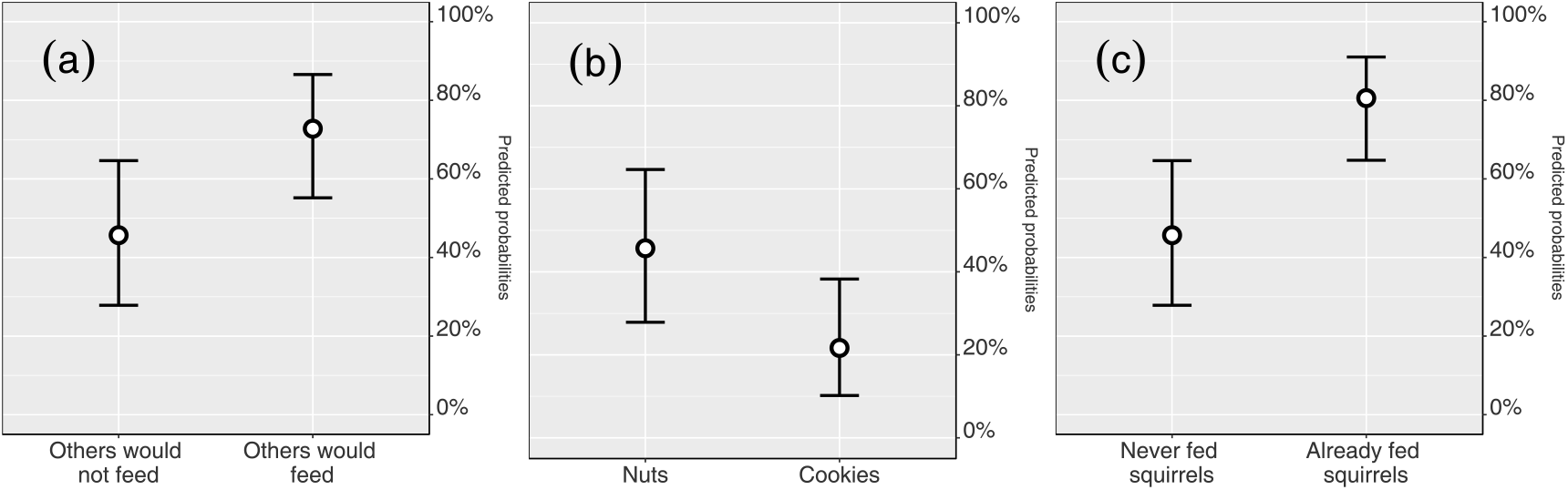
Marginal effects of the model: predicted probabilities for empirical expectations (a), food type (b) and past behavior (c).

## Discussion

To the best of our knowledge, this study contains two novelties. For example, it is the first one exploring drivers of deliberate artificial feeding for invasive squirrels at urban parks. While various studies explored deliberate feeding of various domestic and wildlife species, either in urban contexts (Davey et al. 2019; Jones and Raynolds 2008; Khor et al. 2018) and at protected areas (Milazzo et al. 2015; Newsome and Rodger 2008; Orams 2002; Penteriani et al. 2017; Walpole 2001), none of them focused on invasive alien mammals, nor on squirrels. This gap is noteworthy, because invasive alien squirrels are successful biological invaders worldwide, and urban parks are often their introduction hotspots, representing the context where they establish viable population first, maybe also due to the advantages provided by supplementary feeding (Bertolino et al. 2014; Bertolino, 2009; Merrick et al. 2016). Moreover, this study is the first adopting an operational social norms theory to explore why visitors decide to feed wildlife at urban parks. Although various human dimensions studies measured norm cristallization and norm curves, to explore interactions between humans and wildlife at protected areas (Anderson et al. 2010; Cerri et al. 2019; Vaske and Whittaker 2004; Whittaker 1997), none of them focused on wildlife feeding at urban parks. Indeed, urban parks represent a context where behavioral interventions, such as informative panels, can be relatively easy to enforce and we believe our approach to be potentially important to understand whether and how normative influence can be used to change visitor behavior.

Our findings indicate that supplementary feeding of grey squirrels, in the study area, can be seen as a form of descriptive norm, according to the taxonomy of Bicchieri et al. (2016). Most visitors, among those who were interviewed, believe feeding is practiced because it is enjoyable and because it helps people connecting with nature, but they do not seem to have strong moral beliefs, nor any particular moral expectations from their reference network. On the other hand, either qualitative interviews and factorial surveys indicated that empirical expectations about visitor behavior are an important driver for the decision of feeding squirrels. Their effect over visitor behavior might be important for reducing supplementary feeding, through informative panels indicating that most visitors do not engage in such activity, while visiting the park. We encourage future studies testing these panels, designed also according to different communication frames, to maximize the effectiveness of the message (Miller et al. 2018).

Future studies should also estimate the prevalence of artificial feeding among visitors. In our study area, feeding is practiced on a regular basis, although only by a minority of visitors. However, the situation might be different at other urban parks, or at particular times of the week (e.g. weekends) or with other introduced squirrel species (Bertolino et al. 2004). Feeding estimates would be useful to craft normative messages, and to decide whether providing information about visitors’ behavior is appropriate: whenever squirrels are foraged by most visitors, providing a wrong information would conflict with what people observe, magnifying norm defection and potentially squirrel feeding (Bicchieri and Dimant 2019). Specialized questioning techniques, which protect respondent’s anonymity might be important to obtain unbiased estimates of artificial feeding (Nuno and St. John 2015).

We found surprising that some respondent emphasized the fact that the food provided by visitors might be dangerous to squirrels. This point emerged spontaneously during interviews and it was confirmed by factorial surveys. To the best of our knowledge, this concern had never emerged during any previous study about interactions between squirrels and visitors, nor into any study addressing wildlife feeding at urban parks. Some visitors were genuinely concerned ad the idea that people could give sweets to squirrels, as confirmed by vignette evaluation and some of them even declared that they would have reproved this behavior in public. This concern over food quality suggests that conservationists might design informative panels also emphasizing the risk posed by inadequate food for squirrels, to discourage squirrel feeding. We do not know how widespread the effect of these panels might be and we encourage future studies adopting structured surveys to enhance the generalizability of our findings, which instead aimed to generate new ideas and were based on purposive sampling. While we believe that empirical expectations can affect most different visitors, as they are a major antecedent of interdependent human behavior (Bicchieri and Xiao 2009; Bicchieri 2016), it is plausible that concerns about the effect of food quality over squirrel health could derive from specific beliefs about animal welfare, which are likely to be possessed only by specific segments of visitors (e.g. visitors with mutualistic value orientations, Manfredo et al. 2009). However, in this case, the use of these messages would be even more important, as they might target segments of visitors who are traditionally reluctant to quit feeding wildlife. In this sense, it would be important to see how the two types of information panels would act on different types of visitors, and, in turn, how this differential effect would contribute to reduce supplementary feeding through times.

In this study we explored, by combining qualitative and quantitative methods, supplementary feeding of invasive grey squirrels at urban parks. Based on a taxonomy for collective human behavior, we showed that supplementary feeding is affected by descriptive norms, governed by visitors’ expectations about the behavior of other visitors at the park. This evidence paves the way for further interventions based on normative influence. Moreover, we found that some visitors are also concerned about the effects of supplementary feeding over the health of squirrels. We believe that this second aspect might also be important to design informative panels, emphasizing the risks connected with supplementary feeding.

Our findings show that social norm theory can be important to obtain a better understanding of human-wildlife interactions, and that they can also guide future behavioral interventions aimed at improving them.

## Supporting information

Supplementary dataset and R script

## References

Anderson LE, Manning RE, Valliere WA, Hallo JC (2010) Normative standards for wildlife viewing in parks and protected areas. Hum Dim Wildl 15: 1–15. https://doi.org/10.1080/10871200903360098

Auspurg K, Hinz T (2014) Factorial survey experiments. Sage Publications, Thousand Oaks

Bertolino S (2008) Introduction of the American grey squirrel (*Sciurus carolinensis*) in Europe: a case study in biological invasion. CurrSci India 95: 903–906.

Bertolino, S (2009) Animal trade and non-indigenous species introduction: the world-wide spread of squirrels. Divers Distrib 15: 701–708. https://doi.org/10.1111/j.1472-4642.2009.00574.x

Bertolino S, Mazzoglio PJ, Vaiana M, Currado I (2004) Activity budget and foraging behavior of introduced *Callosciurus finlaysonii* (Rodentia, Sciuridae) in Italy. J Mammal 85: 254–259. https://doi.org/10.1644/BPR-009

Bertolino S, Di Montezemolo NC, Preatoni DG, Wauters LA, Martinoli A (2014) A grey future for Europe: *Sciurus carolinensis* is replacing native red squirrels in Italy. Biol Invasions 16: 53–62. https://doi.org/10.1007/s10530-013-0502-3

Bertolino S, Genovesi P (2003) Spread and attempted eradication of the grey squirrel (*Sciurus carolinensis*) in Italy, and consequences for the red squirrel (*Sciurus vulgaris*) in Eurasia. Biol Conserv 109: 351–358. https://doi.org/10.1016/S0006-3207(02)00161-1

Bicchieri C (2016) Norms in the wild: How to diagnose, measure, and change social norms. Oxford University Press, Oxford.

Bicchieri C, Dimant E (2019) Nudging with care: The risks and benefits of social information. Public Choice, Forthcoming. https://papers.ssrn.com/sol3/papers.cfm?abstract_id=3321394 Accessed 11 August 2019.

Bicchieri C, Noah T (2017) Applying Social Norms Theory in CATS Programming. https://repository.upenn.edu/pennsong/15/ Accessed 11 August 2019

Bicchieri C, Xiao E (2009) Do the right thing: but only if others do so. J Behav Decis Making 22: 191–208. https://doi.org/10.1002/bdm.621

Bürkner PC (2018) Advanced Bayesian multilevel modeling with the R package brms. The R Journal 10: 395–411. doi:10.32614.RJ-2018-017

Cerri J, Martinelli E, Bertolino S (2019) Graphical factorial surveys reveal the acceptability of wildlife observation at protected areas. J Nat Conserv 50. https://doi.org/10.1016/j.jnc.2019.125720

Cox DT, Gaston KJ (2016) Urban bird feeding: connecting people with nature. PloS one 11. https://doi.org/10.1371/journal.pone.0158717

Davey G, Khor MM, Zhao X (2019) Key beliefs underlying public feeding of free-roaming cats in Malaysia and management suggestions. Hum Dim Wildl 24: 1–13. https://doi.org/10.1080/10871209.2018.1522679

Di Febbraro M, Martinoli A, Russo D, Preatoni D, Bertolino S (2016) Modelling the effects of climate change on the risk of invasion by alien squirrels. Hystrix 27: 1–8. doi:10.4404/hystrix-27.1-11776

Di Febbraro M, Menchetti M, Russo D, Ancillotto L, Aloise G, Roscioni F, Preatoni DG, Loy A, Martinoli A, Bertolino S, Mori E (2019) Integrating climate and land-use change scenarios in modelling the future spread of invasive squirrels in Italy. Divers Distrib 25: 644–659. https://doi.org/10.1111/ddi.12890

Dubois S, Fraser D (2013) A framework to evaluate wildlife feeding in research, wildlife management, tourism and recreation. Animals 3: 978–994. https://doi.org/10.3390/ani3040978

Drury R, Homewood K, Randall S (2011) Less is more: the potential of qualitative approaches in conservation research. Anim Conserv 14: 18–24. https://doi.org/10.1111/j.1469-1795.2010.00375.x

Farrow K, Grolleau G, Ibanez L (2017) Social norms and pro-environmental behavior: a review of the evidence. Ecol Econ 140: 1–13. https://doi.org/10.1016/j.ecolecon.2017.04.017

Galbraith JA, Beggs JR, Jones DN, Stanley MC (2015) Supplementary feeding restructures urban bird communities. PNAS 112: 2648–2657. https://doi.org/10.1073/pnas.1501489112

Gelman A, Carlin JB, Stern HS, Dunson DB, Vehtari A, Rubin DB (2013). Bayesian data analysis. CRC press, Boca Raton.

Gelman A, Lee D, Guo J (2015) Stan: A probabilistic programming language for Bayesian inference and optimization. J Educ Behav Stat 40: 530–543. https://doi.org/10.3102/1076998615606113

Gurnell J, Wauters LA, Lurz PPW Tosi G (2004) Alien species and interspecific competition: effects of introduced eastern grey squirrels on red squirrel population dynamics. J Anim Ecol 73: 26–35. https://doi.org/10.1111/j.1365-2656.2004.00791.x

Hauser OP, Gino F, Norton MI (2018) Budging beliefs, nudging behaviour. Mind and Society 17: 15–26. https://doi.org/10.1007/s11299-019-00200-9

Hwang J, Gottdenker NL, Oh DH, Nam HW, Lee H, Chun MS (2018) Disentangling the link between supplemental feeding, population density, and the prevalence of pathogens in urban stray cats. PeerJ, 6: e4988. http://dx.doi.org/10.7717/peerj.4988

Jones DN, Reynolds JS (2008) Feeding birds in our towns and cities: a global research opportunity. J Avian Biol 39: 265–271. https://doi.org/10.1111/j.0908-8857.2008.04271.x

Khor MM, Davey G, Zhao X (2018) Why do people feed free-roaming cats? The role of anticipated regret in an extended theory of planned behavior in Malaysia. Anthrozoös 31: 101–116. https://doi.org/10.1080/08927936.2018.1406204

Larson LR, Cooper CB, Hauber ME (2016) Emotions as drivers of wildlife stewardship behavior: Examining citizen science nest monitors’ responses to invasive house sparrows. Hum Dim Wildl 21: 18–33. https://doi.org/10.1080/10871209.2015.1086933

Lioy S, Mori E, Wauters LA, Bertolino S (2016) Weight operated see-saw feeding hoppers are not selective for red squirrels when greys are present. Mamm Biol 81: 365–371. https://doi.org/10.1016/j.mambio.2016.03.008

Lioy S., Marsan A., Balduzzi A., Wauters L.A., Martinoli A., Bertolino S. (2019) The management of the introduced grey squirrel seen through the eyes of the media. Biol Invasions 21: 3723–3733. https://doi.org/10.1007/s10530-019-02084-9

Loy A, Aloise G, Ancillotto L, Angelici FM, Bertolino S, Capizzi D, Castiglia R, Colangelo P, Contoli L, Cozzi B, Fontaneto D, Lapini L, Maio N, Monaco A, Mori E, Nappi A, Podestà MA, Sarà Loy A, Aloise G, Ancillotto L, Angelici F M, Bertolino S, Capizzi D, Castiglia R, Colangelo P, Contoli L, Cozzi B, Fontaneto D, Lapini L, Maio N, Monaco A, Mori E, Nappi A, Podestà MA, Sarà M, Scandura M, Russo D, Amori G (2019) Mammals of Italy: an annotated checklist. Hystrix, The Italian Journal of Mammalogy 30: 87–106. doi:10.4404/hystrix\T1\textendash00196-2019

Mackay M, Jennings S, van Putten EI, Sibly H, Yamazaki S (2018) When push comes to shove in recreational fishing compliance, think ‘nudge’. Mar Policy 95: 256–266. https://doi.org/10.1016/j.marpol.2018.05.026

Manfredo MJ, Teel TL, Henry KL (2009) Linking society and environment: A multilevel model of shifting wildlife value orientations in the western United States. Soc Sci Quart 90: 407–427.https://doi.org/10.1111/j.1540-6237.2009.00624.x

Merrick MJ, Evans KL, Bertolino S (2016) Urban grey squirrel ecology, associated impacts, and management challenges. In: Shuttleworth C, Lurz PWW, Gurnell J (ed) The Grey Squirrel: Ecology Management of an Invasive Species in Europe. European Squirrel Initiative, Woodbridge, Suffolk UK, pp. 57–77.

Milazzo M, Badalamenti F, Fernandez TV, Chemello R (2005) Effects of fish feeding by snorkellers on the density and size distribution of fishes in a Mediterranean marine protected area. Mar Biol 146: 1213–1222.https://doi.org/10.1007/s00227-004-1527-z

Miller ZD, Freimund W, Metcalf EC, Nickerson N (2018) Targeting your audience: wildlife value orientations and the relevance of messages about bear safety. Hum Dim Wildl 23: 213–226.https://doi.org/10.1080/10871209.2017.1409371

Murray MH, Becker DJ, Hall RJ, Hernandez SM (2016) Wildlife health and supplemental feeding: a review and management recommendations. Biol Conserv 204: 163–174.https://doi.org/10.1016/j.biocon.2016.10.034

Newsome D, Rodger K (2008) To feed or not to feed: A contentious issues in wildlife tourism. In: Lunney D, Munn A, Meikle W (ed) Too close for comfort: contentious issues in human-wildlife encounters. Royal Zoological Society of New South Wales, Mosman, NSW, Australia, pp 255–270

Nuno A, St John FA (2015) How to ask sensitive questions in conservation: A review of specialized questioning techniques. Biol Conserv 189: 5–15.https://doi.org/10.1016/j.biocon.2014.09.047

Orams MB (2002) Feeding wildlife as a tourism attraction: a review of issues and impacts. Tourism Manage 23: 281–293.https://doi.org/10.1016/S0261-5177(01)00080-2

Penteriani V, López-Bao JV, Bettega C, Dalerum F, del Mar Delgado M, Jerina K, Kojola I, Krofel M, Ordiz A (2017) Consequences of brown bear viewing tourism: A review. Biol Conserv 206: 169–180.https://doi.org/10.1016/j.biocon.2016.12.035

Richardson KM, Doerr V, Ebrahimi M, Lovegrove TG, Parker KA (2015) Considering dispersal in reintroduction and restoration planning. In: Armstrong D, Hayward M, Moro D, Seddon P, Advances in reintroduction biology of Australian and New Zealand fauna, CSIRO Publishing, Clayton, pp 59–72.

Rust NA, Abrams A, Challender DW, Chapron G, Ghoddousi A, Glikman JA, Gowan CH, Hughes C, Rastogi A, Said A, Sutton A, Taylor N, Thomas S, Unnikrishan H, Webber AD, Wordingham G, Hill CM (2017) Quantity does not always mean quality: The importance of qualitative social science in conservation research. Soc Nat Resour 30: 1304–1310.https://doi.org/10.1080/08941920.2017.1333661

Schultz PW, Nolan JM, Cialdini RB, Goldstein NJ, Griskevicius V (2018) The constructive, destructive, and reconstructive power of social norms: Reprise. Perspect Psychol Sci 13: 249–254. https://doi.org/10.1177/1745691617693325

Starkey A, delBarco-Trillo J (2019) Supplementary feeding can attract red squirrels (Sciurus vulgaris) to optimal environments. Mamm Biol 94: 134–139.https://doi.org/10.1016/j.mambio.2018.05.004

Walpole MJ (2001) Feeding dragons in Komodo National Park: a tourism tool with conservation complications. Anim Conserv 4: 67–73.https://doi.org/10.1017/S136794300100107X

Whittaker D (1997) Capacity norms on bear viewing platforms. Hum Dim Wildl 2: 37–49.https://doi.org/10.1080/10871209709359093

Vaske J, Whittaker D (2004) Normative approaches to natural resources. Soc Nat Resour: 283–294.

Venturini M, Franzetti B, Genovesi P, Marsan A, Spanò S (2005) Distribuzione e consistenza della popolazione di scoiattolo grigio Sciurus carolinensis Gmelin, 1788 nel levante genovese. Hystrix 16: 53–58.

