## Supplementary dataset and R script for "Artificial feeding of invasive grey squirrels (*Sciurus carolinensis*) at urban parks: a social norms approach to understand its drivers and to guide behavioral interventions": Script_ScoiattoloValentino.R

```
#Loading packages
{
require(gridExtra)
require(stringr)
require(foreign)
require(xlsx)
require(ggplot2)
require(ggthemr)
require(plotly)
require(dplyr)
require(broom)
require(tidyverse)
require(plyr)
require(reshape2)
require(reshape)
require(brms)
require(sjPlot)
require(purrr)
require(tidyr)
require(devtools)
require(tidybayes)
require(ggridges)
require(cowplot)
}

#############################################################
#Interviews
#############################################################
{
#Demographics
df <- read.csv(file="~/Documents/Jacopo/Research_Projects/Ongoing/Scoiattolo_Valentino/Dataset/Valentino_Interviews.csv")
head(df)
nrow(df)
t(colnames(df))
prop.table(table(df$sex))
mean(df$age)
sd(df$age)
prop.table(table(df$edu))
prop.table(table(df$Italian))
prop.table(table(df$Torino))
prop.table(table(df$child))
prop.table(table(df$dog))
df[which(df$frequency_visit_coded==""), "frequency_visit_coded"] <- NA
df$frequency_visit_coded <- factor(df$frequency_visit_coded, levels=c("FirstTime",
                            "MultipleTimes_year", "MultipleTimes_month", "OnceFew_week", "Everyday"))
prop.table(table(df$frequency_visit_coded))

head(df)
#Disapproval
prop.table(table(df$refnet_disapp)) #30.6 % people have someone in their refnet who will disapprove
nrow(df[which(df$refnet_disapp_who_relatives==1),])/nrow(df[which(df$refnet_disapp==1),]) #n=5
nrow(df[which(df$refnet_disapp_who_friends==1),])/nrow(df[which(df$refnet_disapp==1),]) #n=5
nrow(df[which(df$refnet_disapp_who_visitors==1),])/nrow(df[which(df$refnet_disapp==1),]) #n=2
nrow(df[which(df$refnet_disapp_who_others==1),])/nrow(df[which(df$refnet_disapp==1),]) #n=0
nrow(df[which(df$refnet_disapp==1&df$disapp_empexp==1),])/nrow(df[which(df$refnet_disapp==1),]) #90.1%
nrow(df[which(df$refnet_disapp==1&df$disapp_normex==1),])/nrow(df[which(df$refnet_disapp==1),]) #n=5
nrow(df[which(df$refnet_disapp==1&df$disapp_action==1),])/nrow(df[which(df$refnet_disapp==1),]) #n=5

#Approval
prop.table(table(df$refnet_app)) #27.8% respondents have someone who will approve
nrow(df[which(df$refnet_app_who_relatives==1),])/nrow(df[which(df$refnet_app==1),]) #n=5
nrow(df[which(df$refnet_app_who_friends==1),])/nrow(df[which(df$refnet_app==1),]) #n=5
nrow(df[which(df$refnet_app_who_visitors==1),])/nrow(df[which(df$refnet_app==1),]) #n=2
nrow(df[which(df$refnet_app_who_others==1),])/nrow(df[which(df$refnet_app==1),]) #n=0
nrow(df[which(df$refnet_app==1&df$app_empexp==1),])/nrow(df[which(df$refnet_app==1),]) #90.1%
nrow(df[which(df$refnet_app==1&df$app_normex==1),])/nrow(df[which(df$refnet_app==1),]) #n=5
nrow(df[which(df$refnet_app==1&df$app_action==1),])/nrow(df[which(df$refnet_app==1),]) #n=5

#Moral beliefs
prop.table(table(df$feed_moralbel)) #48.6% of respondents had neutral moral beliefs

#Positive consequences
prop.table(table(df$feed_positiveconseq))
prop.table(table(df$fpc_pleasure)) #Pleasure: 38.9%
prop.table(table(df$fpc_connection)) #Connection with nature or animals: 38.9%
prop.table(table(df$fpc_child)) #Children: 27.8%

#Negative consequences of feeding
prop.table(table(df$feed_negativeconseq))
prop.table(table(df$fnc_dangerous)) #Dangerous for squirrels: 5.6%
prop.table(table(df$fnc_behavior)) #Behavior/aggressive: 13.8%
prop.table(table(df$fnc_number)) #Numerical increase: 25.0%
prop.table(table(df$fnc_disease)) #Disease transmission: 11.1%
prop.table(table(df$fnc_diet)) #Wrong diet: 22.2%

#Quit feeding (is it hard?)
prop.table(table(df$quit_feeding_hard)) #71.4%
prop.table(table(df$qfh_habit)) #Habit: 50.0%
prop.table(table(df$qfh_connection)) #Connection with animals and nature: 5.6%
prop.table(table(df$qfh_mind)) #Mentality: 8.3%
prop.table(table(df$qfh_pleasure)) #Pleasure: 2.7%

#Disapproval
prop.table(table(df$disapproval_belief)) #Disapproving people feeding squirrels (25%)
prop.table(table(df$disapproval_action)) #Criticizin people feeding squirrels (13.7%, 8.3% due to food)

t(colnames(df))
}

#############################################################
#Factorial surveys
#############################################################
{
#Loading the dataset
df <- read.csv(file="~/Documents/Jacopo/Research_Projects/Ongoing/Scoiattolo_Valentino/Dataset/DasetFSE_Fixed.csv")
head(df)
nrow(df)
df$respondent <- paste(df$id, df$wave, paste="")
table(df$respondent)
df$food <- factor(df$food, levels=c("Nuts", "Cookies"))
df$season <- factor(df$season, levels=c("Summer", "Winter"))
df[which(df$sex==""), "sex"] <- NA
df$sex <- factor(df$sex, levels=c("M", "F"))
df[which(df$edu==""), "edu"] <- NA
df$edu <- factor(df$edu, levels=c("elementary", "secondary", "high", "degree"))
df[which(df$age==""), "age"] <- NA
df$age <- factor(df$age, levels=c("18-25 years", "26-35 years", "36-45 years", "46-60 years", "Over 60"))
df$resident <- factor(df$resident, levels=c(0,1), labels=c("Not resident", "Resident"))
df$freqvisit <- factor(df$freqvisit)
df[,c(2:6)] <- as.data.frame(lapply(df[,c(2:6)], function(x) as.factor(x)))

#Exploration
df$one <- 1
foo <- aggregate(one ~ respondent + sex + age + edu+ resident + freqvisit, sum, data=df)
head(foo)
round(prop.table(table(foo$sex)), 2)
round(prop.table(table(foo$age)),3)
round(prop.table(table(foo$edu)), 3)
round(prop.table(table(foo$resident)), 3)
round(prop.table(table(foo$freqvisit)), 3)

#Model fitting
str(df)
modA <- brm(outcome ~ food + child + past + visible + othersfeed + empexpfeed + season + resident + (1|respondent),
           family=bernoulli(link = "logit"), chains=4, cores=6, iter=5000, warmup = 1000, data=df)
plot_model(modA) + theme_bw() + labs(title="") +
  geom_hline(aes(yintercept=1), linetype="dashed", size=1.5, color="firebrick")
save(modA, file="~/Documents/Jacopo/Research_Projects/Ongoing/Scoiattolo_Valentino/Dataset/Models/modA.rda")

#No season
modB <- brm(outcome ~ food + child + past + visible + othersfeed + empexpfeed + resident + (1|respondent),
            family=bernoulli(link = "logit"), chains=4, cores=6, iter=5000, warmup = 1000, data=df)
save(modB, file="~/Documents/Jacopo/Research_Projects/Ongoing/Scoiattolo_Valentino/Dataset/Models/modB.rda")

#No child
modC <- brm(outcome ~ food + past + visible + othersfeed + empexpfeed + resident + (1|respondent),
            family=bernoulli(link = "logit"), chains=4, cores=6, iter=5000, warmup = 1000, data=df)
save(modC, file="~/Documents/Jacopo/Research_Projects/Ongoing/Scoiattolo_Valentino/Dataset/Models/modC.rda")

#No others feeding
modD <- brm(outcome ~ food + past + visible + empexpfeed + resident + (1|respondent),
            family=bernoulli(link = "logit"), chains=4, cores=6, iter=5000, warmup = 1000, data=df)
save(modD, file="~/Documents/Jacopo/Research_Projects/Ongoing/Scoiattolo_Valentino/Dataset/Models/modD.rda")

#No resident
modE <- brm(outcome ~ food + past + visible + empexpfeed + (1|respondent),
            family=bernoulli(link = "logit"), chains=4, cores=6, iter=5000, warmup = 1000, data=df)
save(modE, file="~/Documents/Jacopo/Research_Projects/Ongoing/Scoiattolo_Valentino/Dataset/Models/modE.rda")

#No visible
modF <- brm(outcome ~ food + past + empexpfeed + (1|respondent),
            family=bernoulli(link = "logit"), chains=4, cores=6, iter=5000, warmup = 1000, data=df)
save(modF, file="~/Documents/Jacopo/Research_Projects/Ongoing/Scoiattolo_Valentino/Dataset/Models/modF.rda")

#Model selection
wAIC <- waic(modA, modB, modC, modD, modE, modF)
kfoldCV <- kfold(modA, modB, modC, modD, modE, modF)
save(kfoldCV, file="~/Documents/Jacopo/Research_Projects/Ongoing/Scoiattolo_Valentino/Dataset/Models/kfoldCV.rda")

#Loading models
load(file="~/Documents/Jacopo/Research_Projects/Ongoing/Scoiattolo_Valentino/Dataset/Models/modA.rda")
load(file="~/Documents/Jacopo/Research_Projects/Ongoing/Scoiattolo_Valentino/Dataset/Models/modB.rda")
load(file="~/Documents/Jacopo/Research_Projects/Ongoing/Scoiattolo_Valentino/Dataset/Models/modC.rda")
load(file="~/Documents/Jacopo/Research_Projects/Ongoing/Scoiattolo_Valentino/Dataset/Models/modD.rda")
load(file="~/Documents/Jacopo/Research_Projects/Ongoing/Scoiattolo_Valentino/Dataset/Models/modE.rda")
load(file="~/Documents/Jacopo/Research_Projects/Ongoing/Scoiattolo_Valentino/Dataset/Models/modF.rda")
load(file="~/Documents/Jacopo/Research_Projects/Ongoing/Scoiattolo_Valentino/Dataset/Models/kfoldCV.rda")
wAIC <- waic(modA, modB, modC, modD, modE, modF)
kfoldCV
cbind(as.data.frame(wAIC[c(1:6),]), as.data.frame(kfoldCV[c(1:6),]))
?kfold
#Best model
mod <- modF

#Posterior predictive checks
plot(mod)

pp_check(mod)  # Good
bayes_R2(mod)

bayes_R2(modA)
bayes_R2(modB)
bayes_R2(modC)
bayes_R2(modD)
bayes_R2(modE)
bayes_R2(modF)

summary(mod)

str(df)

#Residuals
res <- residuals(mod)
head(res)
res[,1]
a <- res[,1]
b <- df[which(!is.na(df$outcome)),"outcome"]
res.df <- data.frame(a, b)
ggplot(data=res.df, aes(x=a, y=b)) + geom_point() + geom_smooth(formula=y~x, method=lm) #Standard residuals
plot(a) #Randomized quantile residuals

#Marginal effects
lab.idx <- c("Past \nbehavior", "Food: cookies \nvs nuts", "Empirical \nexpectations")
p1 <- plot_model(mod) + theme_bw() + labs(title = "") +
  scale_x_discrete(labels=lab.idx) +
  geom_hline(aes(yintercept=1), linetype="dashed", size=1.5, color="firebrick") +
  theme(axis.text.x = element_text(colour="grey20",size=16,angle=0,hjust=.5,vjust=.95,face="plain"),
        axis.text.y = element_text(colour="grey20",size=16,angle=0,hjust=1,vjust=0,face="plain"),  
        axis.title.x = element_text(colour="grey20",size=16,angle=0,hjust=.5,vjust=0,face="plain"),
        axis.title.y = element_text(colour="grey20",size=16,angle=90,hjust=.5,vjust=.5,face="plain"),
        panel.border = element_rect(fill = NA))
p1

p <- marginal_effects(mod, effect="empexpfeed")
est <- p$empexpfeed$estimate__
lcl <- p$empexpfeed$lower__
ucl <- p$empexpfeed$upper__
value <- c("No", "Yes")
d <- data.frame(est, lcl, ucl, value)
lab.idx <- c("Others would \nnot feed", "Others would \nfeed")
p2 <- ggplot() + geom_errorbar(data=d, mapping=aes(x=value, ymin=lcl, ymax=ucl), width=0.2, size=1) + 
  geom_point(data=d, mapping=aes(x=value, y=est), size=4, shape=21, colour="black", fill="white", stroke=2) +
  labs(x = "", y = "Predicted probabilities") + scale_x_discrete(labels=lab.idx) +
  scale_y_continuous(breaks=c(0, 0.2, 0.4, 0.6, 0.8, 1.0), labels=c("0%", "20%", "40%", "60%", "80%", "100%"),
                     limits=c(0,1), position="right") + theme_gray() +
  theme(axis.text.x = element_text(colour="grey20",size=16,angle=0,hjust=.5,vjust=.95,face="plain"),
        axis.text.y = element_text(colour="grey20",size=16,angle=0,hjust=1,vjust=0,face="plain"),  
        axis.title.x = element_text(colour="grey20",size=16,angle=0,hjust=.5,vjust=0,face="plain"),
        axis.title.y = element_text(colour="grey20",size=12,angle=90,hjust=.5,vjust=.5,face="plain"),
        panel.border = element_rect(fill = NA)) +
  annotate("text", x = 0.6, y = 0.9, size=10, hjust=0, label = "(a)", parse = TRUE)

p <- marginal_effects(mod, effect="food")
est <- p$food$estimate__
lcl <- p$food$lower__
ucl <- p$food$upper__
value <- c("Nuts", "Cookies")
d <- data.frame(est, lcl, ucl, value)
d$value <- factor(d$value, levels=c("Nuts", "Cookies"))
p3 <- ggplot() + geom_errorbar(data=d, mapping=aes(x=value, ymin=lcl, ymax=ucl), width=0.2, size=1) + 
  geom_point(data=d, mapping=aes(x=value, y=est), size=4, shape=21, colour="black", fill="white", stroke=2) +
  labs(x = "", y = "Predicted probabilities") +
  scale_y_continuous(breaks=c(0, 0.2, 0.4, 0.6, 0.8, 1.0), labels=c("0%", "20%", "40%", "60%", "80%", "100%"),
                     limits=c(0,1), position="right") + theme_gray() +
  theme(axis.text.x = element_text(colour="grey20",size=16,angle=0,hjust=.5,vjust=.95,face="plain"),
        axis.text.y = element_text(colour="grey20",size=16,angle=0,hjust=1,vjust=0,face="plain"),  
        axis.title.x = element_text(colour="grey20",size=16,angle=0,hjust=.5,vjust=0,face="plain"),
        axis.title.y = element_text(colour="grey20",size=12,angle=90,hjust=.5,vjust=.5,face="plain"),
        panel.border = element_rect(fill = NA)) +
  annotate("text", x = 0.6, y = 0.9, size=10, hjust=0, label = "(b)", parse = TRUE)


p <- marginal_effects(mod, effect="past")
est <- p$past$estimate__
lcl <- p$past$lower__
ucl <- p$past$upper__
value <- c("No", "Yes")
d <- data.frame(est, lcl, ucl, value)
lab.idx <- c("Never fed \nsquirrels", "Already fed \nsquirrels")
p4 <- ggplot() + geom_errorbar(data=d, mapping=aes(x=value, ymin=lcl, ymax=ucl), width=0.2, size=1) + 
  geom_point(data=d, mapping=aes(x=value, y=est), size=4, shape=21, colour="black", fill="white", stroke=2) +
  labs(x = "", y = "Predicted probabilities") + scale_x_discrete(labels=lab.idx) +
  scale_y_continuous(breaks=c(0, 0.2, 0.4, 0.6, 0.8, 1.0), labels=c("0%", "20%", "40%", "60%", "80%", "100%"),
                     limits=c(0,1), position="right") + theme_gray() +
  theme(axis.text.x = element_text(colour="grey20",size=16,angle=0,hjust=.5,vjust=.95,face="plain"),
        axis.text.y = element_text(colour="grey20",size=16,angle=0,hjust=1,vjust=0,face="plain"),  
        axis.title.x = element_text(colour="grey20",size=16,angle=0,hjust=.5,vjust=0,face="plain"),
        axis.title.y = element_text(colour="grey20",size=12,angle=90,hjust=.5,vjust=.5,face="plain"),
        panel.border = element_rect(fill = NA)) +
  annotate("text", x = 0.6, y = 0.9, size=10, hjust=0, label = "(c)", parse = TRUE)


p1
grid.arrange(p2, p3, p4, nrow=3)

#Coefficients
summary(mod)
est <- c(-0.17, -1.12, 1.61, 1.17)
se <- c(0.39, 0.35, 0.36, 0.36)
lcl <- c(-0.95, -1.83, 0.92, 0.49)
ucl <- c(0.60, -0.45, 2.34, 1.90)
eff.sample <- c(12458, 19964, 20947, 18329)
rhat <- c(rep(1.0, 4))
exp(est)
d <- data.frame(est, se, lcl, ucl, eff.sample, rhat)
names(d) <- c("Est", "S.E", "Lower95%CI", "Upper95%CI", "Eff.Sample", "Rhat")
rownames(d) <- c("Intercept", "Emp. expectations", "Food", "Past behavior")
d[,c(1:4)] <- as.data.frame(lapply(d[,c(1:4)], function(x) exp(x)))
d
}
```
